# Spatio-temporal control of phenylpropanoid biosynthesis by inducible complementation of a cinnamate 4-hydroxylase mutant

**DOI:** 10.1101/2020.07.15.203091

**Authors:** Jeong Im Kim, Christopher Hidalgo-Shrestha, Nicholas D. Bonawitz, Rochus B. Franke, Clint Chapple

## Abstract

Cinnamate 4-hydroxylase (C4H) is a cytochrome P450-dependent monooxygenase that catalyzes the second step of the general phenylpropanoid pathway. Arabidopsis *reduced epidermal fluorescence 3* (*ref3*) mutants, which carry hypomorphic mutations in *C4H*, exhibit global alterations in phenylpropanoid biosynthesis and have developmental abnormalities including dwarfing. Here we report the characterization of a conditional Arabidopsis C4H line (*ref3-2*^*pOpC4H*^), in which wild-type *C4H* is expressed in the *ref3-2* background. Expression of *C4H* in plants with well-developed primary inflorescence stems resulted in restoration of fertility and the production of substantial amounts of lignin, revealing that the developmental window for lignification is remarkably plastic. Following induction of *C4H* expression in *ref3-2*^*pOpC4H*^, we observed rapid and significant reductions in the levels of numerous metabolites, including several benzoyl and cinnamoyl esters and amino acid conjugates. These atypical conjugates were quickly replaced with their sinapoylated equivalents, suggesting that phenolic esters are subjected to substantial amounts of turnover in wild-type plants. Furthermore, using localized application of dexamethasone to *ref3-2*^*pOpC4H*^, we show that phenylpropanoids are not transported appreciably from their site of synthesis. Finally, we identified a defective Casparian strip diffusion barrier in the *ref3-2* mutant root endodermis, which is restored by induction of *C4H* expression.

**Highlight:** The work presented this paper provides evidence of metabolite turnover, plasticity of the developmental window for lignification, and the impact of reduced and restored cinnamate-4-hydroxylase (*C4H)* expression on the Casparian strip.

## INTRODUCTION

Phenylpropanoids are a group of specialized metabolites made mostly from phenylalanine through the phenylpropanoid pathway. Downstream and end products of this pathway play crucial roles in plant growth and stress adaptation. Sinapate esters accumulated in leaf epidermis prevent penetration of ultraviolet (UV)-B radiation into the underlying mesophyll cells where photosynthesis occurs (Ruegger and Chapple, 2001; Dean *et al*., 2014). Flavonoids as anti-oxidants scavenge excess reactive oxygen species which are accumulated during severe/prolonged stress conditions and eventually damage cell organelles (Fini *et al*., 2011; Brunetti *et al*., 2013). Flavonols are also involved in pollen tube development (Muhlemann *et al*., 2018). Monolignols are polymerized to form lignin which provides to plants both the hydrophobicity of vascular bundles and rigidity in general. Another notable site of lignin deposition is the Casparian strip in the root endodermis, which is crucial for water relations and selective uptake of nutrients (Pfister *et al*., 2014).

Biochemical and genetic studies in a broad range of plants have elucidated the biosynthetic pathway of phenylpropanoids (Vogt, 2010; Fraser and Chapple, 2011). The pathway starts with deamination of phenylalanine to produce cinnamic acid (CA) by phenylalanine ammonia lyase (PAL), after which cinnamate 4-hydroxylase (C4H), a cytochrome P450-dependent monooxygenase (CYP73), catalyzes the conversion of CA to *p*-coumaric acid (Koukol and Conn, 1961; Russell and Conn, 1967; Russell, 1971; Schilmiller *et al*., 2009). Thereafter, a series of biochemical reactions produce various structures of phenylpropanoids with differentiated functions (Bonawitz and Chapple, 2010; Fraser and Chapple, 2011). Because the activities of general phenylpropanoid biosynthetic enzymes such as PAL and C4H are required for downstream phenylpropanoid production including lignin, disruption of these enzymes can impact the entire phenylpropanoid pathway. PAL and C4H are regulated through transcriptional, post-translational, and both feedforward and feedback regulation mechanisms (Lamb and Rubery, 1976; Bell-Lelong *et al*., 1997; Jin *et al*., 2000; Olsen *et al*., 2008; Bonawitz *et al*., 2012; Yin *et al*., 2012). PALs are subject to ubiquitination, and polyubiquitinated PALs are degraded by the 26S proteasome (Zhang *et al*., 2013; Yu *et al*., 2019). The interaction of C4H or other CYP enzymes with scaffold proteins forms a lignin metabolon which influences metabolic flux through the different branches of the phenylpropanoid/monolignol biosynthetic pathway (Gou *et al*., 2018).

Given that PAL and C4H function at the first two steps of the general phenylpropanoid pathway, defects in PAL and C4H can reduce flux for phenylpropanoid production including monolignols and flavonoids (Elkind *et al*., 1990; Rohde *et al*., 2004; Schilmiller *et al*., 2009; Huang *et al*., 2010). Although PAL is encoded by a multigene family in the *Arabidopsis thaliana* genome, there is only a single copy of the gene encoding C4H (*C4H*, At2g30490). Disruption of *C4H* is seedling lethal, and Arabidopsis mutants lacking C4H do not form true leaves when grown on soil (Schilmiller *et al*., 2009). A forward genetic screen for mutants with a *reduced epidermal fluorescence* (*ref*) phenotype identified a series of *ref3* mutants possessing leaky *C4H*activity due to mis-sense mutations in C4H (Ruegger and Chapple, 2001; Schilmiller *et al*., 2009). *ref3* mutants show substantial reduction of phenylpropanoids including lignin and sinapate esters. Instead, they accumulate cinnamoylmalate that is not abundant in wild-type extracts (Schilmiller *et al*., 2009). Besides these biochemical abnormalities, *ref3* mutants display unexpected growth defects, such as stem swellings at nodes, abnormal leaf morphology, and dwarfism.

Despite some progress in our understanding of the biochemistry of shoot lignin biosynthesis, that of Casparian strip lignin formation has been advanced only recently. Molecular genetic approaches have identified important factors for the correct localization and assembly of lignin polymerizing scaffolds and their components in Casparian strip formation (Lee *et al*., 2013; Kamiya *et al*., 2015). Pharmacological studies of C4H inhibition indicated the necessity of phenylpropanoid-derived monolignols for a functional Casparian strip barrier. However, conclusive genetic evidence for the requirement of phenylpropanoids in Casparian strip formation is missing since mutants in late lignin biosynthesis genes (CAD, CCR) had only moderate effects on Casparian strip properties (Naseer *et al*., 2012).

To better understand the dynamics of secondary metabolite pools and plasticity of plant growth and lignification, we employed a chemically inducible expression system using *ref3-2* mutants, which allows us to control C4H activity spatially and temporally. Here we report the finding of novel benzoyl and cinnamoyl amino acid conjugates accumulated in Arabidopsis *C4H* mutants, turnover of metabolites upon C4H induction, and the impact of reduced and restored *C4H* expression on Casparian strip.

## MATERIALS and METHODS

### Plant material and growth conditions

*Arabidopsis thaliana* ecotype Col-0 was used as the wild type. The *ref3-2* mutant has been described previously (Schilmiller *et al*., 2009). Plants were grown at 21 ± 1°C under a long-day photoperiod (16h light/ 8h dark) with a light intensity of 100 μE m^−2^ s^−1^.

### Generation of *ref3-2*^*pOpC4H*^ plants and induction of C4H

The Arabidopsis *C4H* open reading frame was amplified by PCR using the primer pair CC2077 and CC2078 (Supplementary Table S1). The PCR product was cloned into a Gateway entry vector pCC1155 which was modified from pENTR1A (Thermo Scientific) (Kim *et al*., 2014). The resulting *C4H* entry clone was recombined with the pOpOn vector (Craft *et al*., 2005) to generate the *pOpON:C4H* construct, which was introduced into *ref3-2* heterozygous plants via *Agrobacterium*-mediated transformation using *A. tumefaciens* strain GV3101. T_1_ transgenic plants were screened on Murashige and Skoog media (Murashige and Skoog, 1962) containing 50 μg mL^-1^ kanamycin. Selected T1 plants showing *ref3-2* morphology were transferred to soil and sprayed with 20 μM dexamethasone (dex), 0.01% Triton X-100 to promote their robust development. The *ref3-2* mutation was detected via PCR amplification with primers CC2396 and CC2397 (Supplementary Table S1) and *Hinf*I digestion as described previously (Schilmiller *et al*., 2009), allowing the establishment of *ref3-2*^*pOpC4H*^ lines homozygous for *ref3-2* and with a single, segregating copy of the *pOpON:C4H* transgene. For continuous induction of C4H, *ref3-2*^*pOpC4H*^ plants were sprayed with 20 μM dex, 0.01% Triton X-100 twice a week.

### Quantitative RT-PCR

Total RNA was isolated from Arabidopsis whole aerial parts using the RNeasy Plant Mini Kit (Qiagen). After DNase treatment (TURBO DNase, ThermoFisher Scientific), complementary DNA was synthesized using a high-capacity cDNA RT kit (ThermoFisher Scientific) following the manufacturer’s protocol. Quantitative RT-PCR was performed using Fast SYBR Green Master Mix (ThermoFisher Scientific). *C4H* was amplified using the primer set of CC3258 and CC3259 (Supplementary Table S1). At1g13320 was used as an internal control with the primer set CC2558 and CC2559 (Supplementary Table S1) used for amplification (Czechowski *et al*., 2005), and relative expression of *C4H* was calculated using the 2^-ΔΔCT^ method (Livak and Schmittgen, 2001).

### Soluble metabolite analysis

Fresh tissues were extracted with 50% (v/v) methanol at 65°C for 1 h. Samples were centrifuged at 16,000 g for 5 min and then samples were analyzed. For HPLC analysis, 10 μL of extract was loaded onto a Shim-pack XR-ODS column (3.0 × 75 mm, 2.2 μm) (Shimadzu) using a gradient from 5% acetonitrile in 0.1% formic acid to 25% acetonitrile in 0.1% formic acid at a flow rate of 1 mL min^-1^. Sinapic acid, cinnamic acid, kaempferol and cyanidin were used as standards for the quantification of sinpoylmalate, cinnamoylmalate, kaempferol glycosides, and anthocyanin, respectively. Liquid chromatography/mass spectrometry (LC/MS) analysis was conducted as following a previous report (Bonawitz *et al*., 2012). Briefly, 10 μL of extract was run on the same Shim-pack XR-ODS column described above with a flow rate of 0.5 mL min^-1^ at an increasing concentration of acetonitrile from 10% to 25% over 14 min and from 25% to 70% over 8 min in 0.1% formic acid. Metabolites were analyzed with an Agilent 1100 LC/MSD TOF mass spectrometer in negative electrospray ionization (ESI) mode.

### Lignin Analysis

Acetylbromide soluble lignin (ABSL) contents were quantified from stems of 8-week-old plants as described previously (Chang *et al*., 2008; Kim *et al*., 2014). Plant samples were ground in liquid nitrogen, washed with 80% ethanol five times, then washed with acetone and dried. Dried samples were incubated in a 4:1 (v/v) mixture of acetic acid/acetyl bromide at 70°C for 2 h. Samples were cooled to room temperature and then transferred to 50-mL volumetric flasks containing 2 M NaOH, acetic acid, and 7.5 M hydroxylamine hydrochloride. Absorbance at 280 nm of each sample was measured and then, absorbance values were converted to mass using the extinction coefficient of 23.29 L g^-1^ cm^-1^. The derivatization followed by reductive cleavage (DFRC) lignin analysis was performed as previously reported (Lu and Ralph, 1998; Weng *et al*., 2010).

### Phloroglucinol-HCl Staining

For histochemical staining, the base of primary inflorescence stems was embedded in 5% agar and cut into 100 μm sections using a vibratome. Sections were then incubated in 16% ethanol (v/v), 10% HCl (v/v), 0.2% phloroglucinol (w/v) for 10 min, then washed, mounted in water, and examined using an Olympus Vanox-S light microscope.

### GUS staining

For GUS staining, fresh plant tissue was incubated in staining solution [50 mM sodium phosphate buffer (pH 7.2), 0.5 mM potassium ferricyanide, 0.5 mM potassium ferrocyanide, 0.3% Triton X-100, and 1.9 mM X-gluc (5-bromo-4-chloro-3-indolyl-β-D-glucuronic acid) at 37°C for 6 h. Then, chlorophyll in the samples was removed with 70% ethanol at 60°C for 4 h, and samples were visualized.

### In vitro plant growth

Seeds were sterilized using 70% ethanol containing 0.05 % Triton X-100. After 10 min shaking at room temperature, the sterilization solution was removed and seeds were washed three times with 96% ethanol before air-drying. Sterilized seeds were placed on half strength Murashige and Skoog media-agar plates, stratified overnight at 4°C in the dark and then placed in the growth chamber with plates oriented vertically. Seven-day-old seedlings with ∼1 cm long primary roots were used for propidium iodide diffusion assays and staining of Casparian strip lignin as described below. For induction and feeding experiments, 1000x stock solutions of 10 mM coniferyl alcohol and 10 mM dex were prepared in DMSO and filter sterilized before being added to autoclaved half strength Murashige and Skoog media-agar.

### Propidium iodide diffusion assays

Vertically grown seedlings were incubated in 15 μM propidium iodide (PI) (Life Technologies, Darmstadt, Germany) for 10 min in the dark. Stained seedlings were rinsed twice in water and subsequently mounted on microscope slides for UV-light microscopy using an Axioplan microscope (Carl Zeiss, Jena, Germany). The stained apoplast was visualized with the filter set: excitation filter BP 500-550 nm, beam splitter FT 580 nm, blocking filter LP 590 nm. Images were recorded with a Nikon DXM-1200 digital camera (Nikon, Düsseldorf, Germany). For the quantitative analysis the point where propidium iodide does not infiltrate the central cylinder was scored and the distance to the “onset of elongation” was determined as number of endodermal cells according to Nasser et al., 2012.

### Casparian strip lignin staining with basic fuchsin

Basic fuchsin staining was performed according to the adapted ClearSee protocol (Ursache *et al*., 2018). Seedlings were fixed in 4% paraformaldehyde in PBS solution for at least 30 min at room temperature. Seedlings were then rinsed twice for 1 min in PBS, transferred to ClearSee solution and incubated overnight, followed by staining in 0.2% basic fuchsin in ClearSee solution overnight. Subsequently, excess basic fuchsin was removed, seedlings were rinsed once in ClearSee solution for 30 min with gentle agitation, followed by overnight washing in fresh ClearSee solution. Finally, seedlings were mounted on slides with ClearSee solution for imaging with both a Leica SP5 confocal laser scanning microscope (Leica, Wetzlar, Germany) and an Olympus Fluoview FV1000 (Olympus, Centre Valley, PA, USA). To image basic fuchsin we used 550 or 561 nm excitation and detected fluorescence at 570-650 nm.

## RESULTS

### Induction of C4H substantially alleviates the metabolic and developmental abnormalities of the *ref3-2* mutant

The Arabidopsis *ref3-2* mutant carries a mis-sense mutation (R249K) in *C4H*, displays abnormal leaf and stem morphology, and is male sterile (Fig. 1A, C) (Schilmiller *et al*., 2009). To explore the impacts of loss and gain of C4H activity on phenylpropanoid metabolism and on plant growth and development, we generated a chemically inducible *C4H* expression system in the *ref3-2* background using a stringent glucocorticoid-inducible vector pOpON, which we call *ref3-2*^*pOpC4H*^ hereafter. pOpON carries a bidirectional inducible promoter and has been shown to effectively control the expression of target genes in Arabidopsis by dex application (Craft *et al*., 2005; Moore *et al*., 2006; Kim *et al*., 2014). Without dex application, *ref3-2* and *ref3-2*^*pOpC4H*^ are indistinguishable (Fig. 1C). Both have small curled epinastic leaves (Fig. 1C), are dwarf and sterile, and produce reduced amounts of sinapoylmalate (Fig. 2A) and flavonoids (Fig. 2B). Without dex application, *C4H* expression in *ref3-2*^*pOpC4H*^ is maintained at wild-type levels, indicating tight control of *C4H* expression in *ref3-2*^*pOpC4H*^ (Fig. 1E). A single application of dex on three-week-old plants increases the expression of *C4H* seven fold within two hours and the induction is maintained at an elevated level for at least four days (Fig. 1E). Based on these results, to fully chemically complement *ref3-2*^*pOpC4H*^, we applied dex twice a week to activate C4H continuously. This treatment substantially restores leaf morphology and plant size (Fig. 1D), fertility, and phenylpropanoid production (Fig. 2) of *ref3-2* mutants, further confirming that a deficiency in C4H activity leads to the perturbations of metabolism and growth in *ref3-2* (Fig. 1, 2).

**Fig. 1.**
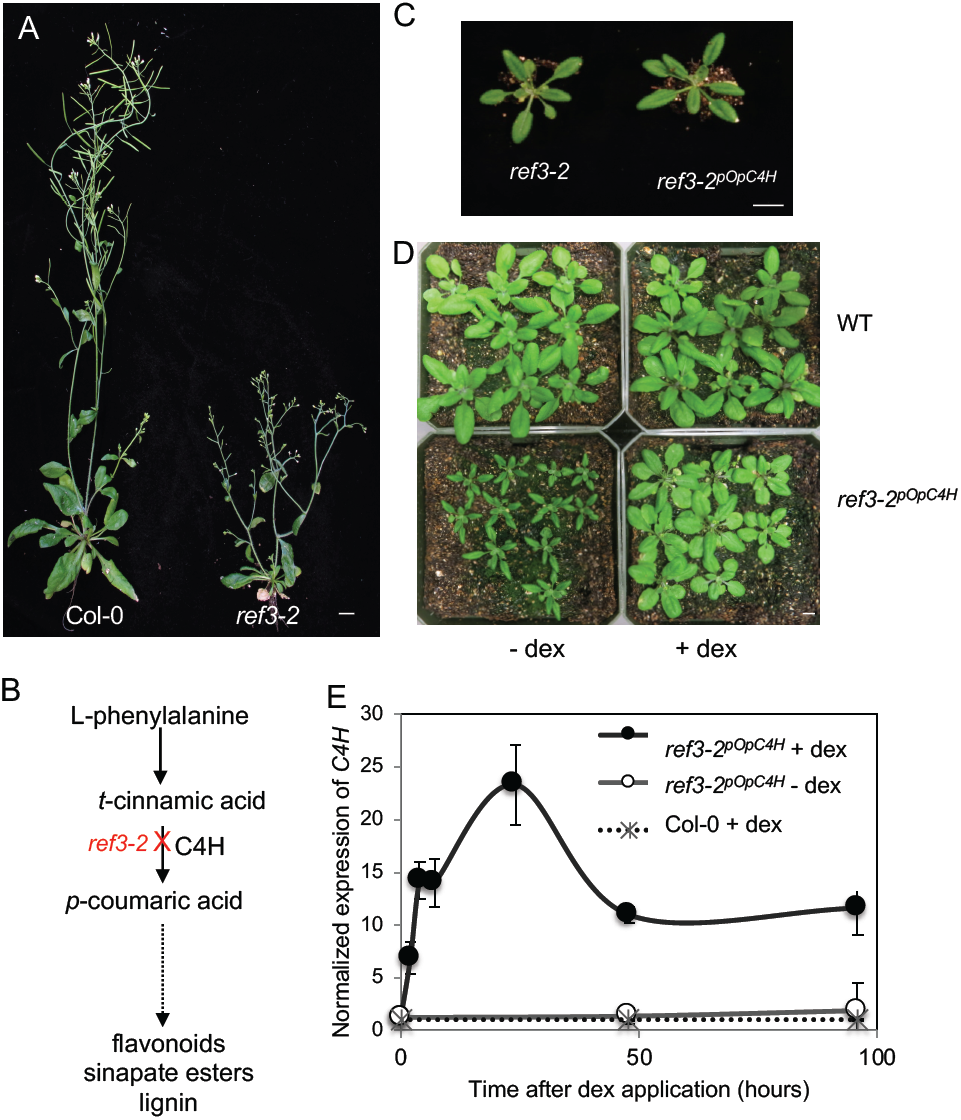
Chemically induced expression of *C4H* rescues the phenotype of *ref3-2* mutant. A. Six-week-old wild-type and *ref3-2* plants. B. *ref3-2* has a defect in C4H the enzyme normally responsible for conversion of *t*-cinnamic aid to *p*-coumaric acid. C. Representative four-week old *ref3-2* and *ref3-2*^*pOpC4H*^ plants. D. *ref3-2*^*pOpC4H*^ (bottom) and wild-type (top) plants grown with dex application (right) or without dex (left) for four weeks. Dex was applied twice a week beginning at 0 days after planting (DAP). E. Relative *C4H* expression in *ref3-2*^*pOpC4H*^ after dex application compared with wild type and untreated *ref3-2*^*pOpC4H*^. Data represent mean ± SD (n=3). Dex was applied one time on three-week-old plants grown on soil and total RNA was extracted from whole rosette leaves. Scale bars = 1 cm.

**Fig. 2.**
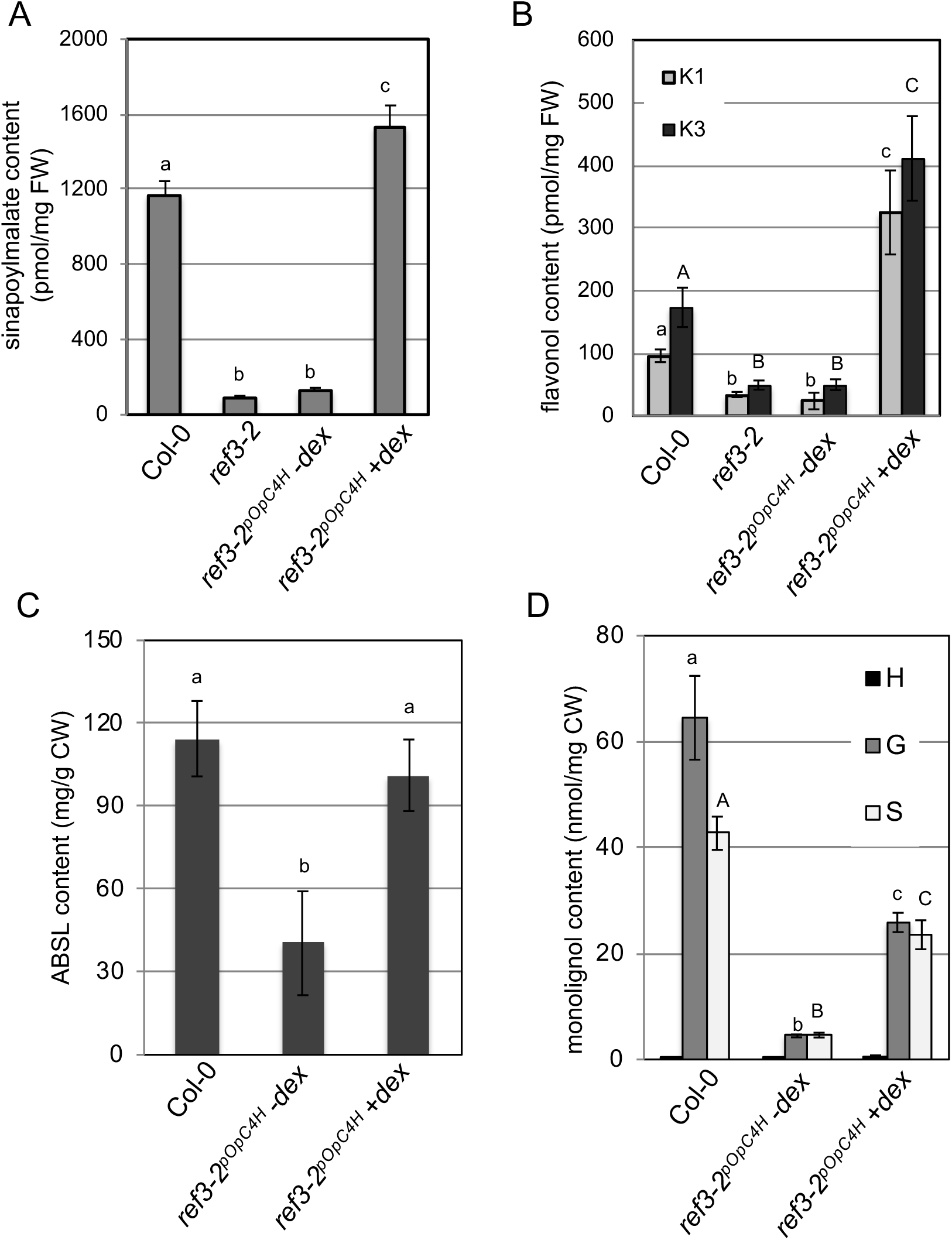
Dex-induced expression of *C4H* restores soluble metabolites and lignification of *ref3-2*^*pOpC4H*^ plants. A. Sinapoylmalate quantification of wild-type and *ref3-2* plants, and of *ref3-2*^*pOpC4H*^ plants either with or without dex application. Dex treatments were performed twice weekly from 0 DAP. B. Flavonol levels of the plants outlined in A. C. Acetylbromide soluble lignin (ABSL) content in stems of 8-week-old wild type and *ref3-2*^*pOpC4H*^ plants with or without dex application. D. DFRC lignin monomer composition of the plants in C. H, *p*-hydroxyphenyl lignin; G, guaiacyl lignin; S, syringyl lignin. Data represent mean ± SD (n=4 for A-C, n=3 for D). The means were compared by one-way ANOVA, and statistically significant differences (P < 0.05) were identified by Tukey’s test and indicated by letters to represent difference between groups. K 1, kaempferol 3-*O*-[6’’-*O*-(rhamnosyl)glucoside] 7-*O*-rhamnoside; K3, kaempferol 3-*O*-rhamnoside 7-*O*-rhamnoside.

Interestingly, the levels of both sinapoylmalate (Fig. 2A) and flavonoids (Fig. 2B) in dex-applied *ref3-2*^*pOpC4H*^ are higher than in wild type. Given that *C4H* is over-expressed upon dex application (Fig. 1E), it is possible that transcriptional regulation of *C4H* limits phenylpropanoid production in wild type. In contrast, acetylbromide soluble lignin (ABSL) content is not higher than the level seen in wild type (Fig. 2C). Wild-type lignin consists of about 60% of guaiacyl-(G-) derived subunits and 40% of syringyl-(S-) derived subunits which gives an S/G ratio of 0.67, whereas *ref3-2* mutants contain reduced amounts of both G and S subunits with an S/G ratio of 1.0 (Fig. 2D). The activation of C4H increases the total amounts of G and S lignin significantly but does not restore the S/G ratio to that of wild-type plants (Fig. 2D). These data may indicate that more complex regulatory mechanisms are involved in the determination of lignin content compared to soluble metabolism, or it may indicate that dex is less effective at reaching lignifying cells than the leaf cells that synthesize soluble phenylpropanoids.

### Temporal control of *C4H* expression reveals rapid turnover of novel phenylpropanoids in *ref3-2* mutant

To determine whether the *ref3-2* mutant phenotypes were irreparable in adult plants, we induced C4H in plants that had already begun flowering. Under the growth conditions employed, one-month-old wild-type plants have fully expanded rosette leaves and primary inflorescence stems, whereas untreated *ref3-2*^*pOpC4H*^ plants have small down-curled leaves and short stems (Fig. 3A). In addition, strong *ref3* alleles give rise to an anthocyanin-deficient phenotype similar to that of *transparent testa* mutants (Koornneef, 1981; Schilmiller *et al*., 2009). When dex was applied to mature *ref3-2*^*pOpC4H*^ plants, accumulation of anthocyanin at the base of stems and petioles was visible as early as two days after C4H induction (Fig. 3A, C). To evaluate the impact of C4H induction on leaf development, we applied dex to *ref3-2*^*pOpC4H*^ plants and examined leaf morphological changes after induction. As shown in Fig. 3C, the small narrow leaves of *ref3-2*^*pOpC4H*^ grow broader and larger upon C4H induction. Thus, despite the appearance that the leaves of mature *ref3* mutants have fully expanded, they in fact retain the ability to expand further.

**Fig. 3.**
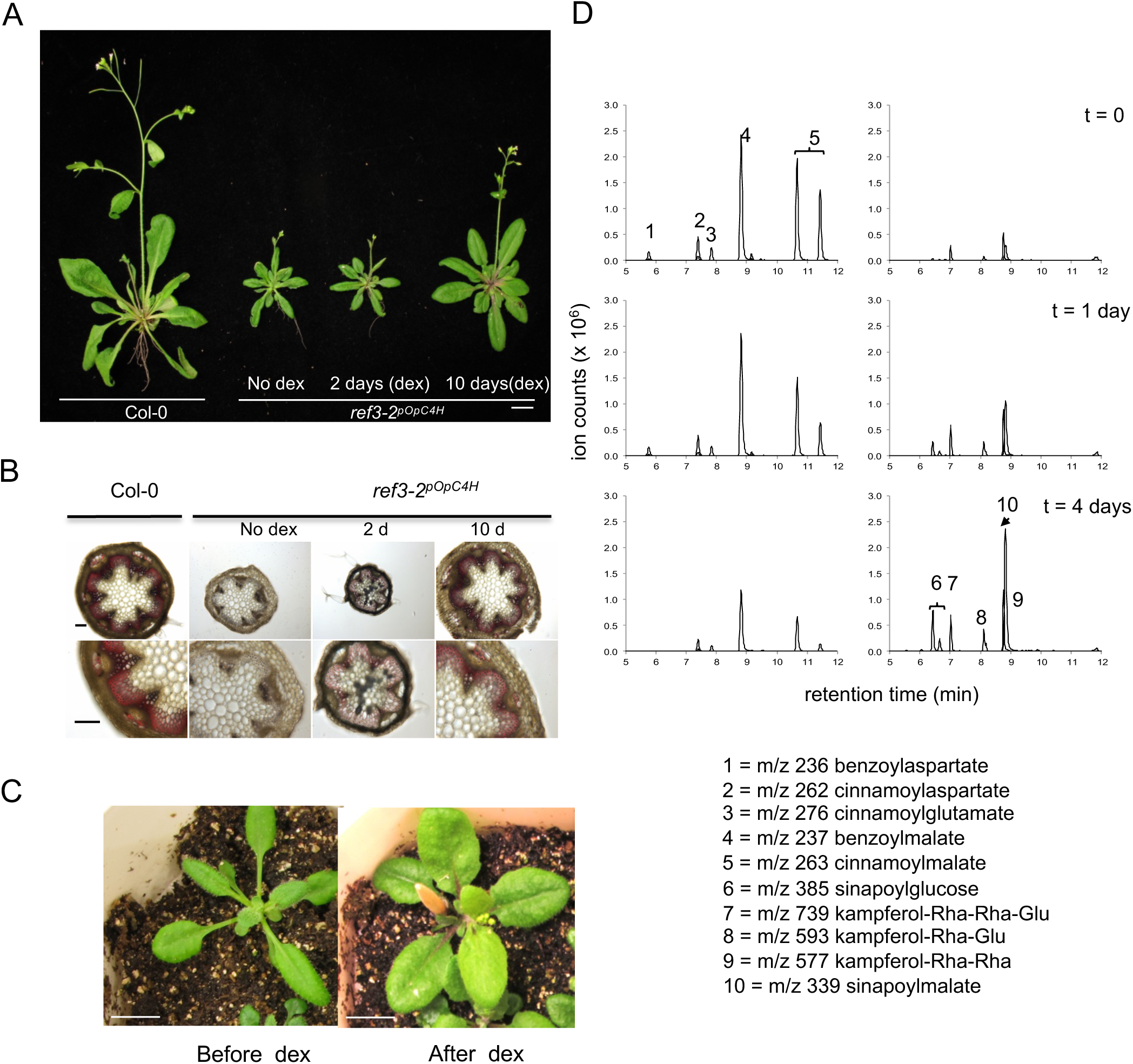
Induction of *C4H* in mature plants leads to restored growth and phenylpropanoid metabolism. A. Forty-day-old wild type and *ref3-2*^*pOpC4H*^. *ref3-2*^*pOpC4H*^ plants were either untreated, or treated with dex for either 2 d or 10 d before being photographed. Scale bar is 1cm. B. Representative stem sections stained with phloroglucinol-HCl. Stem sections were made from primary inflorescences of plants described in A. Scale bars = 100 μm. C. Restored leaf morphology of *ref3-2*^*pOpC4H*^ plants. *ref3-2*^*pOpC4H*^ plants were photographed before and 7 d after dex application. Scale bars = 1cm. D. Induction of *C4H* results in reduced levels of cinnamoyl-and benzoyl-substituted metabolites and increased accumulation of sinapate esters and flavonols. Shown are overlaid extracted ion chromatograms (EICs) corresponding to ten dex/C4H-responsive metabolites as determined immediately preceding (top), 1 d after (middle), or 4 d after (bottom) dex treatment of *ref3-2*^*pOpC4H*^ plants. For clarity, EICs of metabolites whose levels decrease upon *C4H* induction are shown on the left, while those that increase are shown on the right. The m/z values and inferred identities of the labeled peaks are as follows: (1) m/z 236, benzoylaspartate; (2) m/z 262, cinnamoylaspartate; (3) m/z 276, cinnamoylglutamate; (4) m/z 237, benzoylmalate; (5) m/z 263, cinnamoylmalate; (6) m/z 385, sinapoylglucose; (7) m/z 739, kaempferol 3-*O*-[6"-*O*-(rhamnosyl) glucoside] 7-*O*-rhamnoside; (8) m/z 593, kaempferol 3-*O-*glucoside 7-*O-*rhamnoside; (9) m/z 577, kaempferol 3-*O*-rhamnoside 7-*O*-rhamnoside; (10) m/z 339, sinapoylmalate.

To explore whether lignin, once laid down, can be further modified by later C4H induction, we analyzed the lignification status of *ref3-2*^*pOpC4H*^ using a histochemical staining method. Primary inflorescences of wild type and *ref3-2*^*pOpC4H*^ were sectioned and stained with phloroglucinol, a dye that is often used for lignin staining and localization (Kim *et al*., 2014). The weak coloration of stem sections of uninduced *ref3-2*^*pOpC4H*^ indicates lower lignification compared to wild type (Fig. 3B). Ten days after dex application, stem sections of *ref3-2*^*pOpC4H*^ show stronger coloration with phloroglucinol compared with uninduced samples. This indicates that stems of fully matured plants can be lignified even after the normal developmental window for lignification has passed.

To explore the dynamics of secondary metabolite pools, we evaluated metabolite profiles of *ref3-2*^*pOpC4H*^ plants before and after the induction of C4H. In accordance with previous reports, *ref3-2*^*pOpC4H*^ mutants accumulated cinnamoylmalate (peak 5), which wild-type plants do not (Fig. 3D, 4D) (Schilmiller *et al*., 2009). The levels of cinnamoylmalate decreased upon dex application, whereas sinapate esters (peaks 6 and 10) and kaempferol glycosides (peaks 7-9) accumulated (Fig. 3D). In addition to cinnamoylmalate, we found that *ref3-2* mutants accumulate compounds with m/z values and isotopolog ratios consistent with benzoylmalate (peak 4), cinnamoylaspartate (peak 2), benzoylaspartate (peak 1) and cinnamoylglutamate (peak 3). The activation of C4H decreases the benzoyl and cinnamoyl esters and amino acid conjugates in *ref3-2*^*pOpC4H*^ plants (Fig. 3D), indicating that these metabolites are subject to relatively rapid turnover.

### Spatial control of C4H expression showed that phenylpropanoids are not transported appreciably from their site of synthesis

The chemically inducible dex system makes possible the expression of C4H in confined areas. We applied dex on selected portions of *ref3-2*^*pOpC4H*^ leaves and analyzed whether localized complementation of the mutant phenotype in that area affected metabolite profiles in adjacent areas. Because the bidirectional inducible promoter of the pOpC4H vector activates the expression of both C4H and GUS upon dex application, we were able to use β-glucuronidase activity as confirmation of successful dex-mediated transcriptional induction. We first applied dex on leaves and examined them under UV illumination three days after the induction. *ref3-2*^*pOpC4H*^ plants display the characteristic *ref* phenotype, exhibiting reddish fluorescence under UV due to the lack of sinapoylmalate on the leaf epidermis and chlorophyll fluorescence from the mesophyll beneath. Dex application led to localized blue fluorescence, strongly suggesting the accumulation of sinapoylmalate in cells where C4H was activated (Fig. 4B). Consistently, β-glucuronidase activity was only observed where sinapoylmalate accumulated (Fig. 4C). We repeated the same experiments with five independent plants and quantified soluble metabolites in the third and fourth rosette leaves with and without dex application. The dex-treated leaves accumulated more sinapoylmalate and flavonols than untreated leaves (Fig. 4E, F) and cinnamoylmalate was less abundant in dex-treated leaves compared with untreated leaves (Fig. 4D). Similarly, we applied dex on a confined area of *ref3-2*^*pOpC4H*^ stems (Fig. 5A). To repress anthocyanin biosynthesis gene expression, we covered a part of stem with aluminum foil to block light (Meng *et al*., 2004). Anthocyanin accumulation was observed where dex was applied, but only if that tissue was exposed to light. Dex-treated tissue that was shielded from light did not accumulate significant amounts of anthocyanins (Fig. 5A and B). In accordance with a previous report, our data confirm that diffusion or lateral transport of dex in the leaves and stems is limited (Craft *et al*., 2005). These data also indicate that hydroxycinnamate esters and anthocyanins in wild-type and *ref3-2* mutant plants are not transported appreciably from their site of synthesis.

**Fig. 4.**
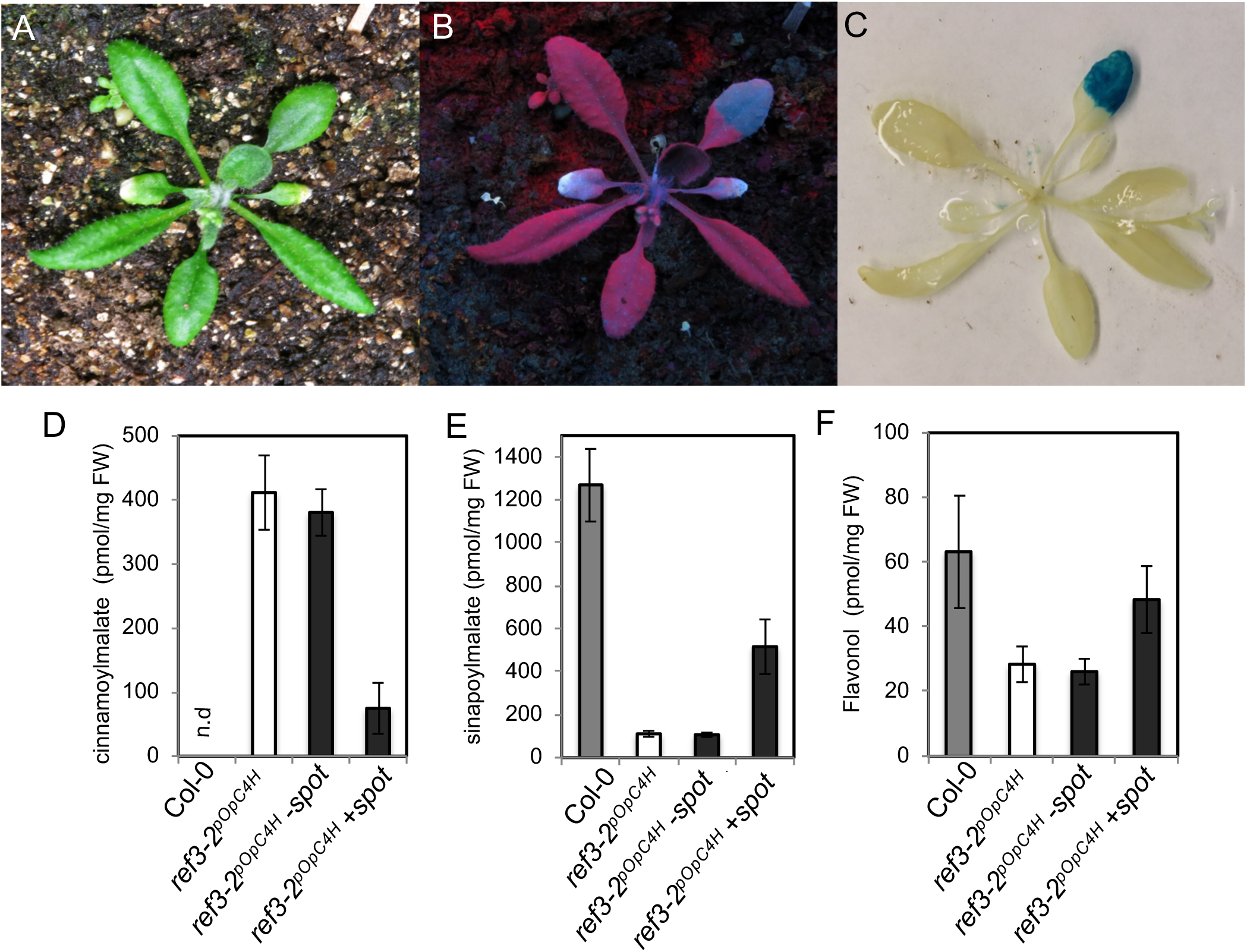
Spatial control of C4H induction in *ref3-2*^*pOpC4H*^ leaves. A-C. Photographs of four-week-old *ref3-2*^*pOpC4H*^ leaves, a small area of which had been treated with dex three days prior, under white light (A), UV light (B) or white light after GUS staining (C). D-F. Quantification of soluble metabolites in dex spot-treated leaves shown in A. Shown are cinnamoylmalate (D), sinapoylmalate (E), and flavonol (F) in water-treated *ref3-2*^*pOpC4H*^ (white bar), water-treated wild-type (gray bar), and water-(spot) or dex treated-leaves (+spot) from the same *ref3-2*^*pOpC4H*^ plants (black bar). Data represent mean ± SD (n=5). n.d.; not detected.

**Fig. 5.**
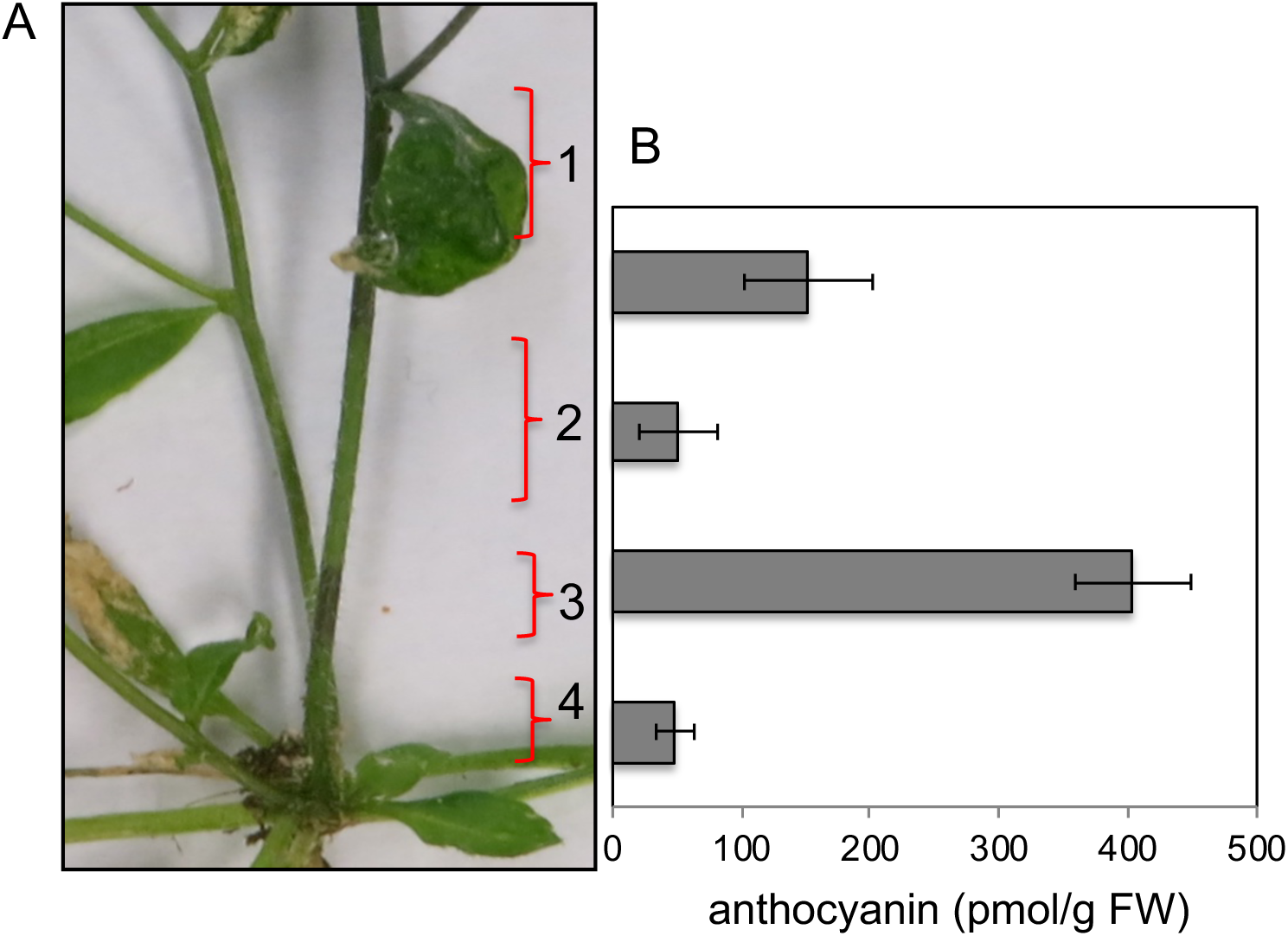
Spatial control of *C4H* induction in *ref3-2*^*pOpC4H*^ stems. A. Segment of a four-week old *ref3-2*^*pOpC4H*^ stem to which dex had been applied at sections 1 to 3. Section 2 of the stem was additionally covered with aluminum foil to block light. Photographs were taken 7 d after dex treatment. B. Anthocyanin (A11, as defined by Tohge *et al*., 2005) content from stem sections described in (A). Data represent mean ± SD (n=4).

### C4H is a key player in the generation of monolignols for functional Casparian strip formation

Previous studies on the spatiotemporal expression pattern of key genes in phenylpropanoid metabolism revealed a strong expression in roots (Bell-Lelong *et al*., 1997; Nair *et al*., 2002). Interestingly, *C4H* expression is not limited to the stele but is also expressed in the endodermis, the tissue in which the Casparian strip develops. This prompted us to investigate a potentially novel phenotype of the *ref3-2* mutant: potentially deficient Casparian strip lignification influenced by reduced C4H activity. Confocal microscopy of basic fuchsin-stained primary roots showed a well-developed, regular Casparian strip network at the 10^th^ endodermal cell after onset of elongation in wild type. In contrast, we observed a non-uniform Casparian strip network in *ref3-2*^*pOpC4H*^ roots at this position with individual endodermal cells apparently lacking Casparian strip lignification (Fig. 6). Moreover, the typically condensed, band-like deposition of Casparian strip lignin is rather diffuse and remains diffuse in distal root zones. This indicates that unhindered carbon flux through the phenylpropanoid pathway is indispensable for the correct and precise deposition of the Casparian strip network.

**Fig. 6.**
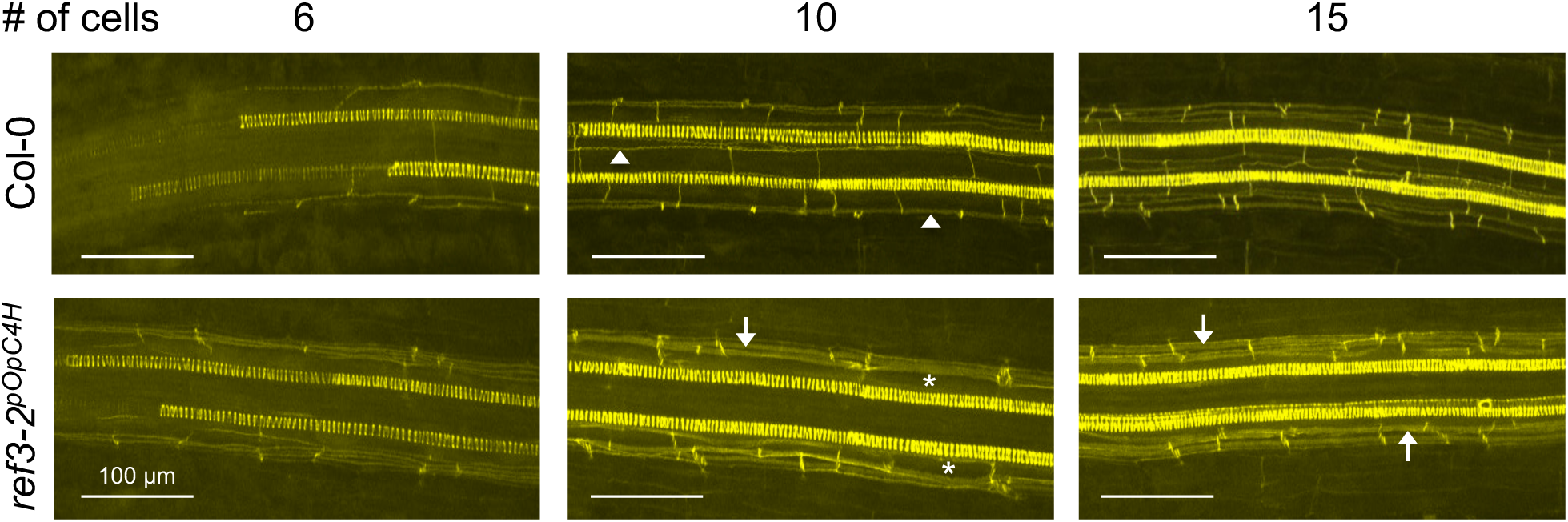
Disruption and partial loss of lignified Casparian strip in *ref3-2*^*pOpC4H*^ roots. Z-stack confocal images of Casparian strip lignin stained with basic fuchsin (yellow). Spiral structures indicate diarchic xylem. In wild-type at about the 10^th^ endodermal cell regular Casparian strip network (arrowheads) is developed. At this stage *ref3-2*^*pOpC4H*^ roots partially lack endodermal lignin (asterisks) or show diffuse lignin deposition (arrows) which is the predominant form in later stage (15^th^ cell). Scale bars = 100 μm.

Although the Casparian strip in the *ref3-2* mutant is diffuse, the hypomorphic *C4H* allele in this genetic background allows some residual lignin to be deposited in the endodermal cell wall. To address the question of whether this residual lignin is sufficient to establish a functional apoplastic barrier, we determined the position were diffusion of an apoplastic tracer (propidium iodide; PI) into the stele is blocked (Fig. 7). In line with previous observations (Naseer *et al*., 2012), in wild-type roots, the Casparian strip diffusion barrier is established at about the 11^th^ endodermal cell after onset of elongation in the endodermis. In contrast, in *ref3-2*^*pOpC4H*^ a PI diffusional barrier is only formed at around the 40^th^ cell, which in these assays corresponds to the root-hypocotyl junction. These data suggest that the endodermis is unable to establish a Casparian strip barrier in the presence of only low levels of C4H. In support of this hypothesis, induction of *C4H* restores the functional endodermal diffusion barrier. Growth of *ref3-2*^*pOpC4H*^ on media containing dex completely restored a wild-type Casparian strip barrier phenotype, indicating that downstream products of C4H are essential for the formation of functional Casparian strips. Furthermore, the exogenously applied monolignol coniferyl alcohol largely complements the deficiency of phenylpropanoid derived Casparian strip precursors in *ref3-2*^*pOpC4H*^. *ref3-2*^*pOpC4H*^ plants grown on media containing 10 µM coniferyl alcohol established a PI diffusion barrier at the 15^th^ cell. Together with the apparent absence of transport of phenylpropanoids among tissues, this indicates that a C4H-controlled biosynthesis of monolignols within the endodermis is vital for construction of the lignin built Casparian strip apoplastic barrier.

**Fig. 7.**
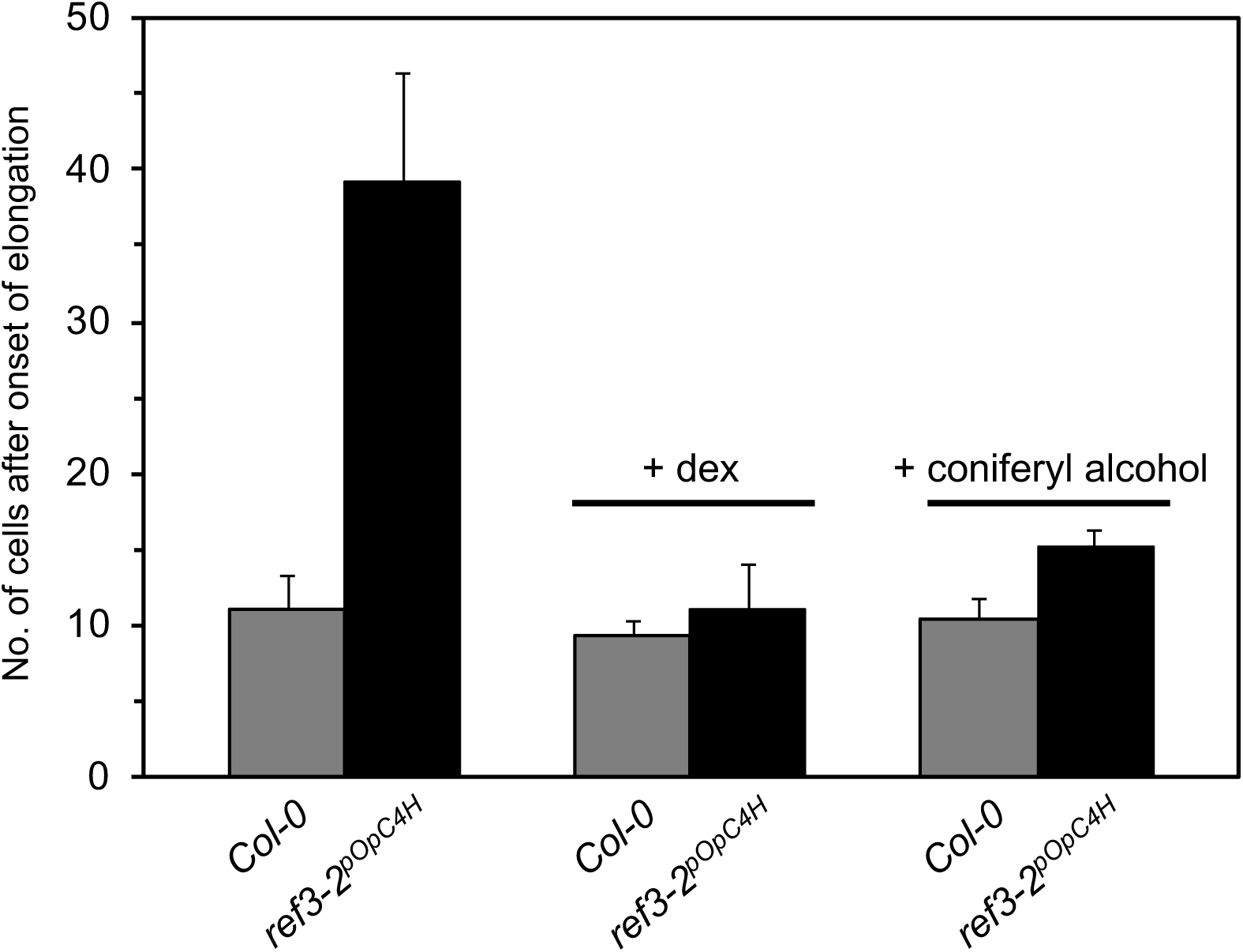
Induction of C4H in primary roots restores monolignol biosynthesis for Casparian strip diffusion barrier. Establishment of a functional apoplastic diffusion barrier determined as the number of endodermal cells at which propidium iodide is blocked at the endodermis. Dex (10 µM) and coniferyl alcohol (10 µM) were applied in Murashige and Skoog medium. Data represent means ± SD (n = 10).

## DISCUSSION

Because C4H functions at an early step of the phenylpropanoid pathway, *ref3*/*c4h* mutants or *C4H* RNAi lines often display pleiotropic growth or biochemical phenotypes (Koopmann *et al*., 1999; Blount *et al*., 2000; Blee *et al*., 2001; Schilmiller *et al*., 2009; Kumar *et al*., 2016). In this study, we investigated the impact of loss or gain of C4H activity on plant growth and phenylpropanoid metabolism through spatiotemporal control of *C4H* expression. The work presented here provides evidence of metabolite turnover, plasticity of the developmental window for lignification, and a role of C4H in Casparian strip formation.

In accordance with previous studies, uninduced *ref3-2*^*pOpC4H*^ plants show a reduction of sinapate esters, flavonoids, and lignin, and the accumulation of cinnamoyl and benzoyl malate, indicative of reduced C4H activity (Fig. 2, 3) (Schilmiller *et al*., 2009; Vanholme *et al*., 2012; Sundin *et al*., 2014). When wild-type C4H is induced, even when the plant has fully matured, the alterations of phenylpropanoid metabolism, including lignification, can be restored (Fig. 3, 4). The decrease in the concentration of benzoyl and cinnamoyl amino acid conjugates upon C4H induction indicates their rapid turnover, possibly including hydrolytic release of benzoate and cinnamate. It is possible that cinnamate is routed into phenylpropanoid synthesis following release via C4H-catalyzed hydroxylation. The accumulation of benzoate derivatives in *ref3* are likely explained by chain shortening of the cinnamic acid that accumulates in the mutant. Their disappearance following *C4H* induction may indicate that they are routed towards the production of benzoylated or benzoate-derived compounds. Benzoate is a precursor of various compounds including benzoylated glucosinolates in Arabidopsis (Lee *et al*., 2012; Widhalm and Dudareva, 2015). Another possible explanation is that the formation of benzoyl and cinnamoyl aspartate/glutamate conjugates may be a part of degradation process like that seen in hormone homeostasis. For example, the levels of free indole-3-acetic acid (IAA), an important auxin, are tightly regulated. Conjugation of IAA is to either glutamate (IAA-Glu) or aspartate (IAA-Asp), abolishes its auxin activity. IAA-Glu and IAA-Asp are then subject to irreversible oxidation, followed by catabolism of IAA, whereas IAA-alanine or -leucine conjugates are storage forms (Normanly, 1997; Ljung *et al*., 2002; Staswick *et al*., 2005). Similarly, benzoyl and cinnamoyl glutamate/aspartate conjugates may be forms of cinnamate and benzoate targeted for degradation. Although the biosynthetic routes and the fates of these compounds remain elusive, it is evident that cinnamate and benzoate conjugates are subject to rapid turnover.

Because the inducible promoter used in this study contains 35S promoter elements, dex application leads to *C4H* over-expression. The fact that dex-treated *ref3-2*^*pOpC4H*^ plants over-produce sinapoylmalate and flavonoids compared to wild type (Fig 2A, B) suggests that transcriptional regulation of *C4H* may limit phenylpropaniod production in wild type. Interestingly, acetylbromide soluble lignin content does not increase to greater than wild-type levels (Fig 2C), and lignin composition is not completely rescued (Fig 2D) under the same C4H activation condition. The high S/G ratio of lignin monomers in dex-treated *ref3-2*^*pOpC4H*^ plants is reminiscent of lignin phenotypes of *ref3-3*, a weak allele which accumulates wild-type levels of Klason lignin and exhibits a high S/G ratio due to lower amounts of G subunits (Schilmiller *et al*., 2009). It appears that the applied dex is insufficient to fully activate C4H in the lignifying cells whereas it activates C4H efficiently in epidermal cells where soluble metabolites are produced. In contrast to studies indicating that lignin precursors can be trafficked between cells (Smith *et al*., 2017), our dex spot testing results indicate that phenylpropanoids may be synthesized in a cell-autonomous manner and are not transported appreciably to neighboring cells in the epidermis.

*ref3-2* mutants show developmental abnormalities including narrow, downcurled leaves, loss of apical dominance and swellings at the base of lateral branches, similar to that of *ref3-1* mutants, which possess another strong *ref3* allele (Fig. 1B, 3) (Schilmiller *et al*., 2009). The cause for this developmental perturbation remains unknown; however, the activation of C4H restores these developmental defects. Of particular interest is the observation that application of dex to mature *ref3-2*^*pOpC4H*^ plants leads to additional leaf expansion (Fig. 3C). This observation indicates that phenylpropanoid products may have a previously undescribed role in leaf expansion. *PAL* quadruple mutants (*pal1/2/3/4*) are dwarf but they do not show this type of abnormal leaf morphology (Huang *et al*., 2010). This suggests that some of developmental abnormality of *ref3* mutants may result from the accumulation of its substrates or their derivatives rather than a deficiency in downstream products. Studies have shown that *cis*-cinnamic acid (*cis*-CA) has growth regulator-like activities (Yang *et al*., 1999; Yin *et al*., 2003). *trans-* cinnamic acid (*trans-*CA) the substrate of C4H, can be photoisomerized to become *cis*-cinnamic acid (*cis*-CA) and is detected in trace amounts in plants (Yin *et al*., 2003; Wong *et al*., 2005). *cis*-CA inhibits the gravitropism of tomato seedlings root growth of Arabidopsis, and auxin transport (Steenackers *et al*., 2017), and it promotes elongation of tomato cells (Yang *et al*., 1999; Yin *et al*., 2003; Wong *et al*., 2005). On the other hand, *trans-*CA and its derivatives enhance auxin signaling and promote leaf expansion in both auxin-dependent and auxin-independent manners (Kurepa *et al*., 2018; Kurepa and Smalle, 2019). Thus, it is possible that an imbalance of these molecules may contribute to abnormal cell growth of C4H-deficient leaves. The identity of molecules or mechanisms leading to the characteristic leaf morphology of *ref3* mutants needs further investigation.

Previously characterized Casparian strip mutants such as *casp1 casp3, myb36* and *esb1* eventually establish a PI diffusion barrier in the endodermis of more mature basal root zones (Hosmani *et al*., 2013; Kamiya *et al*., 2015). However, the primary root endodermis of *ref3-2*, like in *sgn3-3* (Pfister *et al*., 2014), does not block the apoplastic tracer from diffusion into the stele along the entire seedling root, indicating a severe functional Casparian strip defect (Fig 7). Remarkably, *ref3-2* exhibits an extreme growth phenotype compared to other Casparian strip defective mutants that show either only mild (*sng3-3*) or no obvious growth phenotypes (*casp1 casp3, myb36, esb1*) under laboratory conditions. This indicates that *ref3* dwarfism is not caused by limitations in the function of root barriers affecting water use efficiency and/or nutrient homeostasis. Instead, it suggests that phenylpropanoids are not only required for structural polymer biosynthesis but down-stream products of C4H are also involved in plant growth and development.

Basic fuchsin staining revealed an only weak lignin signal in endodermal cell walls (Fig 6). However, root lignification is not completely abolished in the leaky *ref3-2* mutant background. The residual C4H activity appears sufficient to maintain early root xylem development as indicated by an equal basic fuchsin staining in the protoxylem of wild type and *ref3-2*. Although histochemical detection is only semi-quantitative, the lignin staining indicates that spiral cell wall thickenings in the protoxylem represent a major sink for lignin monomers. Alternatively, the strong effect on Casparian strip lignification compared to xylem lignification may reflect tissue-specific *C4H* expression levels that are moderate and become rate limiting more rapidly in the endodermis than in the xylem (Nair *et al*., 2002).

Interestingly, cell wall lignification in the endodermis of *ref3-2* is surprisingly diffuse compared to the precisely localized and condensed lignin deposition forming the Casparian strip network in wild type. This is markedly different from previously characterized Casparian mutants. In *esb1, casp1 casp3, rbohf/sgn4, myb36*, and *sgn3* ectopic lignin accumulates in endodermal cell corners and the central Casparian strip lignin ring is disrupted (Lee *et al*., 2013; Hosmani *et al*., 2013; Pfister *et al*., 2014; Kamiya *et al*., 2015). These mutants exhibit perturbations in lignin polymerization, regulation, or the localization and assembly of the protein scaffolds required for Casparian strip formation. However, they are assumed to have intact phenylpropanoid lignin monomer biosynthesis. The absence of holes in the *ref3-2* Casparian strip and the reestablished PI-diffusion barrier upon external coniferyl alcohol supplementation (Fig. 7) suggests that the apoplastic Casparian strip formation machinery in *ref3-2* is functioning and correctly localized. Still, the diffuse and less restricted lignification in *ref3-2* indicates that phenylpropanoids also contribute to the spatial control and presumably anchoring and condensing of Casparian strip lignin.

The inducible *C4H* expression system in the *ref3-2* background represents a sophisticated tool to also study physiological responses associated with Casparian strips in a detailed and spatiotemporal manner. In addition to restoring fertility to *ref3-2*^*pOpC4H*^ plants such that homozygous mutant seed can be obtained, it enables temporal control of Casparian strip formation. In combination with split agar plates, the restricted mobility of phenylpropanoids would enable the generation of root zones with different developmental stages of Casparian strip formation that can be exploited in water and ion transport studies.

The barrier properties of the Casparian strip are expected to also depend on the chemical composition and structure of the lignin polymer. To date, the small and fragile nature of Arabidopsis roots have hampered the isolation of pure Casparian strips for detailed chemical and structural analysis of the lignin polymer. The complementation of the *ref3-2* Casparian strip defect by coniferyl alcohol is in line with a guaiacyl dominated composition that has been deduced from the analysis of root sections with genetically and hormone-controlled delay in root xylem formation (Naseer *et al*., 2012). It will be interesting to investigate the capacity of other phenylpropanoids to complement the Casparian strip defects in *ref3-2* to evaluate the plasticity of Casparianstrip biosynthesis and composition.

## Supplementary data

Table S1. The list of primers used in this study.

## Data Availability

All data supporting the findings of this study are available within the paper and within its supplementary materials published online.

## Acknowledgements

This work was supported as part of the Center for Direct Catalytic Conversion of Biomass to Biofuels, an Energy Frontier Research Center funded by the US Department of Energy, Office of Science, Basic Energy Sciences under award no. DE-SC0000997. We thank Dr. Jing-Ke Weng for the generation of the C4H entry vector.

## Author Contributions

J.I.K., R.B.F. and C.C. designed the research project; J.I.K., C.H.-S. and N.D.B. performed the experiments and analyzed the data; J.I.K., R.B.F. and C.C. wrote the manuscript.

## Abbreviations

ABSL: acetylbromide soluble lignin
CA: cinnamic acid
C4H: Cinnamate-4-hydroxylase
CW: cell wall
CYP: cytochrome P450-dependent monooxygenase
dex: dexamethasone
DFRC: derivatization followed by reductive cleavage
FW: fresh weight
PAL: phenylalanine ammonia lyase
PI: propidium iodide

## REFERENCES

Bell-Lelong DA, Cusumano JC, Meyer K, Chapple C. 1997. Cinnamate-4-hydroxylase expression in Arabidopsis. Regulation in response to development and the environment. Plant Physiology 113, 729–738.

Blee K, Choi JW, O’Connell AP, Jupe SC, Schuch W, Lewis NG, Bolwell GP. 2001. Antisense and sense expression of cDNA coding for CYP73A15, a class II cinnamate 4-hydroxylase, leads to a delayed and reduced production of lignin in tobacco. Phytochemistry 57, 1159–1166.

Blount JW, Korth KL, Masoud SA, Rasmussen S, Lamb C, Dixon RA. 2000. Altering Expression of Cinnamic Acid 4-Hydroxylase in Transgenic Plants Provides Evidence for a Feedback Loop at the Entry Point into the Phenylpropanoid Pathway. Plant Physiology 122, 107–116.

Bonawitz ND, Chapple C. 2010. The Genetics of Lignin Biosynthesis: Connecting Genotype to Phenotype. Annual Review of Genetics 44, 337–363.

Bonawitz ND, Soltau WL, Blatchley MR, Powers BL, Hurlock AK, Seals LA, Weng J-K, Stout J, Chapple C. 2012. REF4 and RFR1, Subunits of the Transcriptional Coregulatory Complex Mediator, Are Required for Phenylpropanoid Homeostasis in Arabidopsis. Journal of Biological Chemistry 287, 5434–5445.

Brunetti C, Di Ferdinando M, Fini A, Pollastri S, Tattini M. 2013. Flavonoids as Antioxidants and Developmental Regulators: Relative Significance in Plants and Humans. International Journal of Molecular Sciences 14, 3540–3555.

Chang XF, Chandra R, Berleth T, Beatson RP. 2008. Rapid, Microscale, Acetyl Bromide-Based Method for High-Throughput Determination of Lignin Content in Arabidopsis thaliana. Journal of Agricultural and Food Chemistry 56, 6825–6834.

Craft J, Samalova M, Baroux C, Townley H, Martinez A, Jepson I, Tsiantis M, Moore I. 2005. New pOp/LhG4 vectors for stringent glucocorticoid-dependent transgene expression in Arabidopsis. The Plant Journal: For Cell and Molecular Biology 41, 899–918.

Czechowski T, Stitt M, Altmann T, Udvardi MK, Scheible W-R. 2005. Genome-Wide Identification and Testing of Superior Reference Genes for Transcript Normalization in Arabidopsis. Plant Physiology 139, 5–17.

Dean JC, Kusaka R, Walsh PS, Allais F, Zwier TS. 2014. Plant Sunscreens in the UV-B: Ultraviolet Spectroscopy of Jet-Cooled Sinapoyl Malate, Sinapic Acid, and Sinapate Ester Derivatives. Journal of the American Chemical Society 136, 14780–14795.

Elkind Y, Edwards R, Mavandad M, Hedrick SA, Ribak O, Dixon RA, Lamb CJ. 1990. Abnormal plant development and down-regulation of phenylpropanoid biosynthesis in transgenic tobacco containing a heterologous phenylalanine ammonia-lyase gene. Proceedings of the National Academy of Sciences of the United States of America 87, 9057–9061.

Fini A, Brunetti C, Di Ferdinando M, Ferrini F, Tattini M. 2011. Stress-induced flavonoid biosynthesis and the antioxidant machinery of plants. Plant Signaling & Behavior 6, 709–711.

Fraser CM, Chapple C. 2011. The Phenylpropanoid Pathway in Arabidopsis. The Arabidopsis Book 9, e0152.

Gou M, Ran X, Martin DW, Liu C-J. 2018. The scaffold proteins of lignin biosynthetic cytochrome P450 enzymes. Nature Plants 4, 299.

Hosmani PS, Kamiya T, Danku J, Naseer S, Geldner N, Guerinot ML, Salt DE. 2013. Dirigent domain-containing protein is part of the machinery required for formation of the lignin-based Casparian strip in the root. Proceedings of the National Academy of Sciences 110, 14498–14503.

Huang J, Gu M, Lai Z, Fan B, Shi K, Zhou Y-H, Yu J-Q, Chen Z. 2010. Functional Analysis of the Arabidopsis PAL Gene Family in Plant Growth, Development, and Response to Environmental Stress. Plant Physiology 153, 1526–1538.

Jin H, Cominelli E, Bailey P, Parr A, Mehrtens F, Jones J, Tonelli C, Weisshaar B, Martin C. 2000. Transcriptional repression by AtMYB4 controls production of UV-protecting sunscreens in Arabidopsis. The EMBO Journal 19, 6150–6161.

Kamiya T, Borghi M, Wang P, Danku JMC, Kalmbach L, Hosmani PS, Naseer S, Fujiwara T, Geldner N, Salt DE. 2015. The MYB36 transcription factor orchestrates Casparian strip formation. Proceedings of the National Academy of Sciences 112, 10533–10538.

Kim JI, Ciesielski PN, Donohoe BS, Chapple C, Li X. 2014. Chemically Induced Conditional Rescue of the Reduced Epidermal Fluorescence8 Mutant of Arabidopsis Reveals Rapid Restoration of Growth and Selective Turnover of Secondary Metabolite Pools. Plant Physiology 164, 584–595.

Koopmann E, Logemann E, Hahlbrock K. 1999. Regulation and Functional Expression of Cinnamate 4-Hydroxylase from Parsley. Plant Physiology 119, 49–56.

Koornneef M. 1981. The complex syndrome of ttg mutants. Arabidoopsis Information Service 18, 45–51.

Koukol J, Conn EE. 1961. The metabolism of aromatic compounds in higher plants. IV. Purification and properties of the phenylalanine deaminase of Hordeum vulgare. The Journal of Biological Chemistry 236, 2692–2698.

Kumar R, Vashisth D, Misra A, Akhtar MQ, Jalil SU, Shanker K, Gupta MM, Rout PK, Gupta AK, Shasany AK. 2016. RNAi down-regulation of cinnamate-4-hydroxylase increases artemisinin biosynthesis in Artemisia annua. Scientific Reports 6, 26458.

Kurepa J, Shull TE, Karunadasa SS, Smalle JA. 2018. Modulation of auxin and cytokinin responses by early steps of the phenylpropanoid pathway. BMC Plant Biology 18, 278.

Kurepa J, Smalle JA. 2019. trans-Cinnamic acid-induced leaf expansion involves an auxin-independent component. Communicative & Integrative Biology 12, 78–81.

Lamb CJ, Rubery PH. 1976. Phenylalanine ammonia-lyase and cinnamic acid 4-hydroxylase: Product repression of the level of enzyme activity in potato tuber discs. Planta 130, 283–290.

Lee S, Kaminaga Y, Cooper B, Pichersky E, Dudareva N, Chapple C. 2012. Benzoylation and sinapoylation of glucosinolate R-groups in Arabidopsis. The Plant Journal 72, 411–422.

Lee Y, Rubio MC, Alassimone J, Geldner N. 2013. A Mechanism for Localized Lignin Deposition in the Endodermis. Cell 153, 402–412.

Livak KJ, Schmittgen TD. 2001. Analysis of relative gene expression data using real-time quantitative PCR and the 2(-Delta Delta C(T)) Method. Methods (San Diego, Calif.) 25, 402–408.

Ljung K, Hull AK, Kowalczyk M, Marchant A, Celenza J, Cohen JD, Sandberg G. 2002. Biosynthesis, conjugation, catabolism and homeostasis of indole-3-acetic acid in Arabidopsis thaliana. In: Perrot-Rechenmann C,, In: Hagen G, eds. Auxin Molecular Biology. Dordrecht: Springer Netherlands, 249–272.

Lu F, Ralph J. 1998. The DFRC Method for Lignin Analysis. 2. Monomers from Isolated Lignins. Journal of Agricultural and Food Chemistry 46, 547–552.

Meng X, Xing T, Wang X. 2004. The role of light in the regulation of anthocyanin accumulation in Gerbera hybrida. Plant Growth Regulation 44, 243.

Moore I, Samalova M, Kurup S. 2006. Transactivated and chemically inducible gene expression in plants. The Plant Journal 45, 651–683.

Muhlemann JK, Younts TLB, Muday GK. 2018. Flavonols control pollen tube growth and integrity by regulating ROS homeostasis during high-temperature stress. Proceedings of the National Academy of Sciences 115, E11188–E11197.

Murashige T, Skoog F. 1962. A Revised Medium for Rapid Growth and Bio Assays with Tobacco Tissue Cultures. Physiologia Plantarum 15, 473–497.

Nair RB, Xia Q, Kartha CJ, Kurylo E, Hirji RN, Datla R, Selvaraj G. 2002. Arabidopsis CYP98A3 mediating aromatic 3-hydroxylation. Developmental regulation of the gene, and expression in yeast. Plant Physiology 130, 210–220.

Naseer S, Lee Y, Lapierre C, Franke R, Nawrath C, Geldner N. 2012. Casparian strip diffusion barrier in Arabidopsis is made of a lignin polymer without suberin. Proceedings of the National Academy of Sciences of the United States of America 109, 10101–10106.

Normanly J. 1997. Auxin metabolism. Physiologia Plantarum 100, 431–442.

Olsen KM, Lea US, Slimestad R, Verheul M, Lillo C. 2008. Differential expression of four Arabidopsis PAL genes; PAL1 and PAL2 have functional specialization in abiotic environmental-triggered flavonoid synthesis. Journal of Plant Physiology 165, 1491–1499.

Pfister A, Barberon M, Alassimone J, et al. 2014. A receptor-like kinase mutant with absent endodermal diffusion barrier displays selective nutrient homeostasis defects (MJ Harrison, Ed.). eLife 3, e03115.

Rohde A, Morreel K, Ralph J, et al. 2004. Molecular phenotyping of the pal1 and pal2 mutants of Arabidopsis thaliana reveals far-reaching consequences on phenylpropanoid, amino acid, and carbohydrate metabolism. The Plant Cell 16, 2749–2771.

Ruegger M, Chapple C. 2001. Mutations That Reduce Sinapoylmalate Accumulation in Arabidopsis thaliana Define Loci With Diverse Roles in Phenylpropanoid Metabolism. Genetics 159, 1741–1749.

Russell DW. 1971. The Metabolism of Aromatic Compounds in Higher Plants X. PROPERTIES OF THE CINNAMIC ACID 4-HYDROXYLASE OF PEA SEEDLINGS AND SOME ASPECTS OF ITS METABOLIC AND DEVELOPMENTAL CONTROL. Journal of Biological Chemistry 246, 3870–3878.

Russell DW, Conn EE. 1967. The cinnamic acid 4-hydroxylase of pea seedlings. Archives of Biochemistry and Biophysics 122, 256–258.

Schilmiller AL, Stout J, Weng J-K, Humphreys J, Ruegger MO, Chapple C. 2009. Mutations in the cinnamate 4-hydroxylase gene impact metabolism, growth and development in Arabidopsis. The Plant Journal 60, 771–782.

Smith RA, Schuetz M, Karlen SD, Bird D, Tokunaga N, Sato Y, Mansfield SD, Ralph J, Samuels AL. 2017. Defining the Diverse Cell Populations Contributing to Lignification in Arabidopsis Stems. Plant Physiology 174, 1028–1036.

Staswick PE, Serban B, Rowe M, Tiryaki I, Maldonado MT, Maldonado MC, Suza W. 2005. Characterization of an Arabidopsis Enzyme Family That Conjugates Amino Acids to Indole-3-Acetic Acid. The Plant Cell 17, 616–627.

Steenackers W, Klíma P, Quareshy M, et al. 2017. cis-Cinnamic Acid Is a Novel, Natural Auxin Efflux Inhibitor That Promotes Lateral Root Formation. Plant Physiology 173, 552–565.

Sundin L, Vanholme R, Geerinck J, Goeminne G, Höfer R, Kim H, Ralph J, Boerjan W. 2014. Mutation of the Inducible ARABIDOPSIS THALIANA CYTOCHROME P450 REDUCTASE2 Alters Lignin Composition and Improves Saccharification. Plant Physiology 166, 1956–1971.

Tohge T, Nishiyama Y, Hirai MY, et al. 2005. Functional genomics by integrated analysis of metabolome and transcriptome of Arabidopsis plants over-expressing an MYB transcription factor. The Plant Journal 42, 218–235.

Ursache R, Andersen TG, Marhavý P, Geldner N. 2018. A protocol for combining fluorescent proteins with histological stains for diverse cell wall components. The Plant Journal: For Cell and Molecular Biology 93, 399–412.

Vanholme R, Storme V, Vanholme B, Sundin L, Christensen JH, Goeminne G, Halpin C, Rohde A, Morreel K, Boerjan W. 2012. A Systems Biology View of Responses to Lignin Biosynthesis Perturbations in Arabidopsis. The Plant Cell 24, 3506–3529.

Vogt T. 2010. Phenylpropanoid biosynthesis. Molecular Plant 3, 2–20.

Weng J-K, Mo H, Chapple C. 2010. Over-expression of F5H in COMT-deficient Arabidopsis leads to enrichment of an unusual lignin and disruption of pollen wall formation. The Plant Journal: For Cell and Molecular Biology 64, 898–911.

Widhalm JR, Dudareva N. 2015. A Familiar Ring to It: Biosynthesis of Plant Benzoic Acids. Molecular Plant 8, 83–97.

Wong WS, Guo D, Wang XL, Yin ZQ, Xia B, Li N. 2005. Study of cis-cinnamic acid in Arabidopsis thaliana. Plant physiology and biochemistry: PPB 43, 929–937.

Yang XX, Choi HW, Yang SF, Li N. 1999. A UV-light activated cinnamic acid isomer regulates plant growth and gravitropism via an ethylene receptor-independent pathway. Australian Journal of Plant Physiology 26, 325–335.

Yin R, Messner B, Faus-Kessler T, Hoffmann T, Schwab W, Hajirezaei M-R, von Saint Paul V, Heller W, Schäffner AR. 2012. Feedback inhibition of the general phenylpropanoid and flavonol biosynthetic pathways upon a compromised flavonol-3-O-glycosylation. Journal of Experimental Botany 63, 2465–2478.

Yin Z, Wong W, Ye W, Li N. 2003. Biologically active cis-cinnamic acid occurs naturally in Brassica parachinensis. Chinese Science Bulletin 48, 555.

Yu S, Kim H, Yun D-J, Suh MC, Lee B. 2019. Post-translational and transcriptional regulation of phenylpropanoid biosynthesis pathway by Kelch repeat F-box protein SAGL1. Plant Molecular Biology 99, 135–148.

Zhang X, Gou M, Liu C-J. 2013. Arabidopsis Kelch Repeat F-Box Proteins Regulate Phenylpropanoid Biosynthesis via Controlling the Turnover of Phenylalanine Ammonia-Lyase. The Plant Cell 25, 4994–5010.

