## Supplementary material for "Spatio-temporal control of phenylpropanoid biosynthesis by inducible complementation of a cinnamate 4-hydroxylase mutant": Table S1

Supplementary data

Table S1. The list of primers used in this study.

| Name | Sequences 5' to 3' |
|---|---|
| CC2077 | GGGGACAAGTTTGTACAAAAAAGCAGGCTATGGACCTCCTCTGCTGGAG |
| CC2078 | GGGGACCACTTTGTACAAGAAAGCTGGGTTTAACAGTTCCTTGGTTTCAT |
| CC2396 | TTCCGTATCATGTTCGATAG |
| CC2397 | AATGTCAATTTCCCAAAATC |
| CC3258 | TCTCCTCGTGCCTCACATGA |
| CC3259 | TGCTTTCTGCTGGGATATCGTA |
| CC2558 | TAACGTGGCCAAAATGATGCCC |
| CC2559 | GTTCTCCACAACCGCTTGGT |
